# Selective Permeability of Carboxysome Shell Pores to Anionic Molecules

**DOI:** 10.1101/367714

**Authors:** Paween Mahinthichaichan, Dylan M. Morris, Yi Wang, Grant J. Jensen, Emad Tajkhorshid

**Author notes:** Corresponding authors: G.J.J.; E.T. Equal contribution (Paween Mahinthichaichan, Dylan M. Morris, Grant J. Jensen, Emad Tajkhorshid).

## Abstract

Carboxysomes are closed polyhedral cellular microcompartments that increase the efficiency of carbon fixation in autotrophic bacteria. Carboxysome shells consist of small proteins that form hexameric units with semi-permeable central pores containing binding sites for anions. This feature is thought to selectively allow access to RuBisCO enzymes inside the carboxysome by 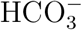 (the dominant form of CO_2_ in the aqueous solution at pH 7.4) but not O_2_, which leads to a non-productive reaction. To test this hypothesis, here we use molecular dynamics simulations to characterize the energetics and permeability of CO_2_, O_2_, and 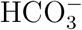 through the central pores of two different shell proteins, namely, CsoS1A of α–carboxysome and CcmK4 of β-carboxysome shells. We find that the central pores are in fact selectively permeable to anions such as 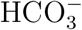, as predicted by the model.

## Introduction

All life depends on the ability of cells to fix atmospheric carbon into organic matter. The key enzyme in this process is ribulose-1,5-bisphosphate carboxylase/oxygenase (RuBisCO), which catalyzes the fixation reaction of CO_2_ and ribulose-1,5-bisphosphate (RuBP) molecules to produce two molecules of 3-phosphoglycerate (3PGA), a precursor molecule for sugar and amino acid biosynthesis. Besides fixing CO_2_ and RuBP, the enzyme fixes O_2_ and RuBP, producing one molecule of 3PGA and one molecule of 2-phosphoglycolate, a wasteful compound [1–5]. RuBisCO is notoriously inefficient with *K_m_* of >150 *µ*M for CO_2_ and *k_cat_* of the reaction in the order of 10 s^*−1*^ [6–8]. It is important to note that the concentration of O_2_ in the atmosphere is ∼21% whereas that of CO_2_ is only ∼0.04%, which is lower than the *K_m_*of CO_2_ for the RuBisCO enzymes [3,5].

To increase the efficiency of RuBisCO, cyanobacteria and carbon-fixing chemoautotrophic bacteria encapsulate RuBisCO and carbonic anhydrase in specialized protein-enclosed cytoplasmic microcompartments called carboxysome [9–12]. To mitigate the occurrence of the O_2_fixation reaction, the carboxysome needs to minimize the penetration of O_2_ into its lumen by mechanisms which would also impact effective entry of CO_2_ due to its chemical resemblance to O_2_. CO_2_ is envisioned to enter the carboxysomal lumen in the form of 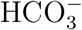 [10, 11, 13], its predominant form at physiological pH. Carbonic anhydrase then converts the 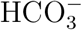 to CO_2_ [14], providing a mechanism for concentrating CO_2_ in the immediate vicinity of the RuBisCO enzymes.

Carboxysomes are classified *α*into *β*and types, based on their composition and evolutionary history [3]. In their native forms, both types resemble icosahedral capsids with a diameter of ∼1,000 Å [9, 15–18]. The outer shell of the carboxysome is formed by the assembly of thousands of copies of a few proteins [2]. *α* carboxysomes are found in *Prochlorococcus*and *Synechococcus* species such as *Halothiobacillus neapolitanus*, and in some other chemoau totrophic bacteria [2, 3, 11, 13, 19]. Their main shell protein is CsoS1A, of which the structure was determined from *H. neapolitanus* [20]. *β* carboxysomes are found in freshwater species such as *Synechococcus elongates PCC 7942* and *Synechocytis sp. PCC 6803* [3, 11, 13, 19], and their main shell proteins are CcmK1–4. These main shell proteins form hexamers and are arranged in a hexagonal lattice with aqueous-exposed surfaces on either side [20–23] (Fig. 1). Along the 6-fold symmetry axis of each hexamer is a pore, termed “the central pore”, with the bottleneck radius of ∼2 Å [2], potentially permitting small molecular species, such as 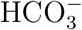, CO_2_ and/or O_2_ molecules to pass through.

**Figure 1:**
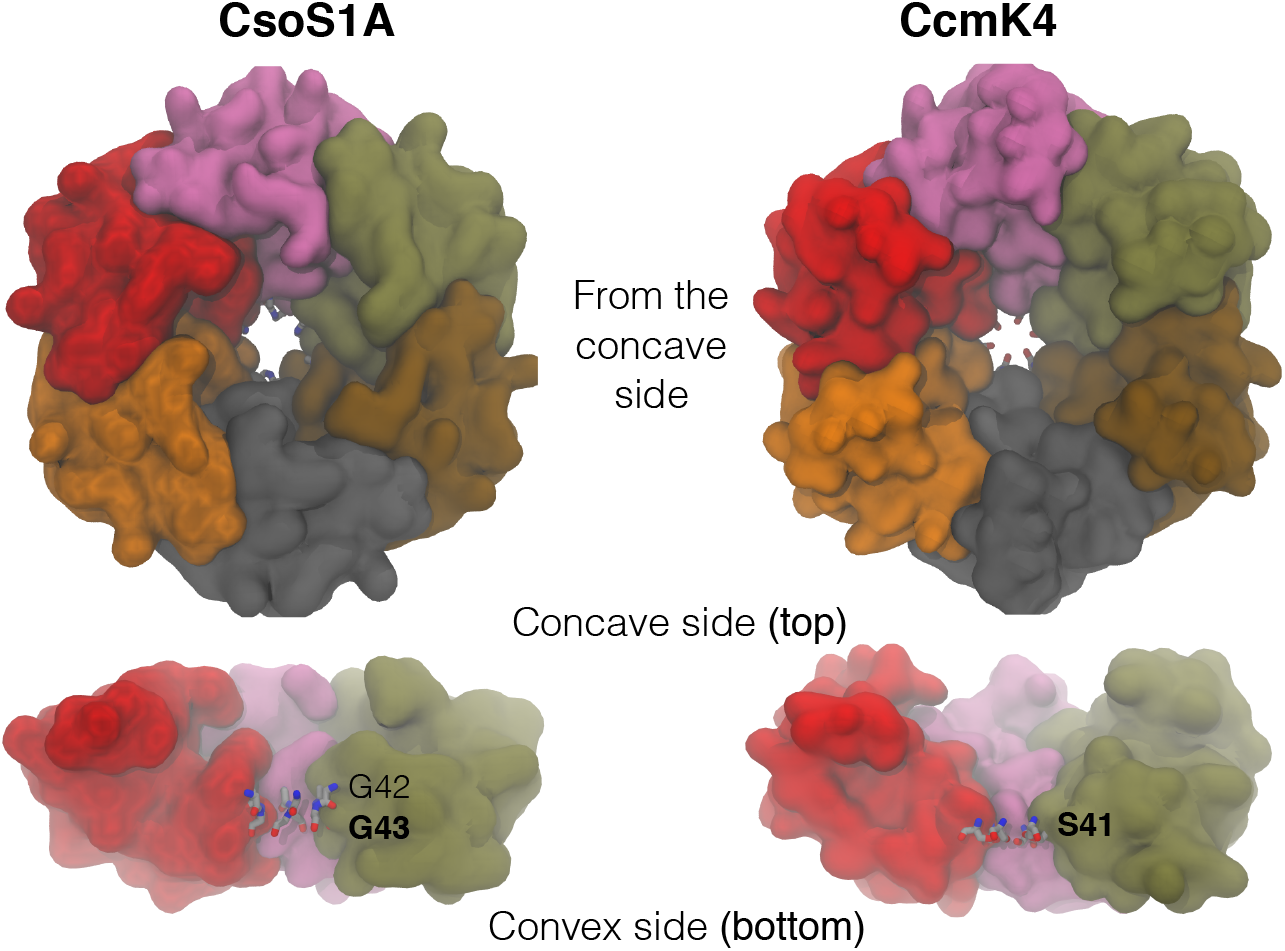
Arrangement of carboxysome shell proteins CsoS1A and CcmK4. Top view from the concave side (upper panel) and side view (lower panel). Individual colored molecular surfaces represent individual subunits of the homohexamers. Each hexamer forms a central pore. The narrowest section (bottleneck) of the pore is formed by six G43 residues for CsoS1A and six S41 residues for CcmK4.

Here, we examine the permeability of 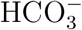, CO_2_, and O_2_ molecules through the central pores of CsoS1A and CcmK4 complexes in full atomic details using molecular dynamics simulations and free energy calculations. The umbrella sampling (US) technique is employed to calculate the free energy profiles for 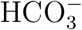, CO_2_, and O_2_ insertion. We find that the central pore of carboxysome shells are preferentially selective for 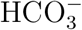, over CO_2_ and O_2_.

## Materials and Methods

### Simulation systems

The simulations were prepared using the crystal structures of CsoS1A from *H. neapolitanus* (PDB entry 2EWH) resolved at 1.4 Å [20] and CcmK4 from *Synechocytis sp. PCC 6803* (PDB entry 2A18) resolved at 2.28 Å [21]. CsoS1A is missing its first five amino acids. CcmK4 is missing its first three amino acids and its last thirteen amino acids. Since these amino acids are located neither near the central pore or at the interface of individual monomeric subunits, they were not modeled. The asymmetric unit of CsoS1A structure is provided by PDB as a monomer while that of CcmK4 structure is provided as a trimer. The biologically relevant hexameric complexes were constructed by VMD [24] using the transformation matrices provided in the PDB files. Each modeled hexamer was centered at the origin so that the 6-fold symmetry axis (and the central pore) coincided with the z-axis. A hexameric periodic box was constructed according to the crystallographic dimensions given in the PDB files. Internal water molecules were added to each complex with DOWSER [25]. The complex was then solvated with TIP3P waters [26] and ionized with 150 mM NaCl. The resulting hydrated hexameric CsoS1A and CcmK4 systems comprised 28,115 and 27,619 atoms, respectively.

### Simulation protocols

The simulations were performed using NAMD2 [27] with a time step of 2 fs and the CHARMM36 force field [28, 29]. The periodic boundary condition (PBC) was used throughout the simulations. All covalent bonds involving hydrogen atoms were kept rigid using the SHAKE algorithm [30]. To evaluate long-range electrostatic interactions in PBC without truncation, the particle mesh Ewald method [31] with a grid density of 1/Å^3^ was used. The cutoff for van der Waals interactions was set at 12 Å. The simulations were performed under NPT ensemble. The temperature was maintained at 300 K by Langevin dynamics [32] with a damping coefficient *γ* of 1 /ps. The Nosé-Hoover Langevin piston method [32, 33] with a piston period of 200 fs was used to maintain the pressure at 1 atm.

### Equilibration and steered molecular dynamics

The equilibrations of the hydrated complexes of CsoS1A and CcmK4 began with 5,000 steps of energy minimization using the conjugated gradient algorithm, followed by 0.5-ns protein heavy-atom restrained, 0.5-ns protein backbone-atom restrained and 1-ns protein C_*α*_-atom restrained simulations with *k* = 1 kcal/mol/Å^2^, and finally 10–20 ns unrestrained simulations. In the CsoS1A system, one of the simulated Cl^−^ ions entered the central pore and bound to residues G43 (of the hexamer) located at the center of the pore during the unrestrained simulation. The binding of a Cl^−^ ion was also observed in the equivalent section of CcmK4, corresponding to the backbones of residues S41. To simulate 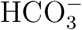, CO_2_ or O_2_ molecule, the bound Cl^−^ molecule was replaced by 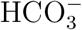, CO_2_ or O_2_ molecules. The force field parameters of these substrate molecules are available in the CHARMM general force field [34]. For the simulation systems with a CO_2_ or O_2_ molecule, one Na^+^ ion was removed for neutralization. 5,000 steps of energy minimization and 100 ps of equilibration were performed on these three systems. Steered molecular dynamics simulations [35] were performed to generate starting structures for the US simulations, described in the following section. The backbone nitrogen atoms of residues G42 of CsoS1A or residues S41 of CcmK4 were used to mark the center of the pore, and these were considered to be at z = 0. Using the center of the pore as the starting point, the localized substrate molecule was pulled out of the pore at a velocity of 10 Å/ns using a force constant *k* = 10 kcal/mol/Å^2^. The simulation consisted of two sets. In one set, the molecule was pulled from z = 0 to z = 20 Å (towards the concave surface). In the other set, it was pulled from z = 0 to z = –15 Å (towards the convex surface). To prevent protein translation artificially induced by the force applied on the pulling of the substrate molecule, the positions of C_*α*_ atoms of residue G6 of CsoS1A and those of residues E11 of CcmK4, located away from the pore, were restrained with *k* =10 kcal/mol/Å

### Umbrella sampling and free energy calculations

The free energy (ΔG) profiles of 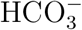, CO_2_ and O_2_ translocation along the central pores were calculated using the US technique [36–38]. The starting frames for the US simulations were taken from the steered molecular dynamics trajectories described above. US simulations spanned from z = –15 Å to z = 20 Å, comprising 71 0.5-Å windows, and each simulation lasted 2.5 ns. This 35-Å length is about the thickness of the carboxysome shell determined from cryo-electron tomography and atomic force microscopy [15, 16, 39]. A harmonic potential with *k* = 10 kcal/mol/Å^2^ was applied to confine the substrate molecule to the center of each window. To construct the ΔG profiles, the last 2-ns trajectories of all of the simulations of a substrate were combined and analyzed using the weighted histogram analysis method (WHAM) [40], with a 0.25-Å histogram bin. Insertion ΔG values of the substrate in individual bins (ΔG_*i*_) were subtracted by ΔG in the bulk solution (ΔG_*bulk*_), yielding relative insertion free energies (ΔG*i,bulk*). The WHAM code was implemented by Professor Alan Grossfield at the University of Rochester Medical Center (http://membrane.urmc.rochester.edu/content/wham).

## Results and Discussion

### Hydration along the central pores

The hydrated complexes of CsoS1A and CcmK4 were simulated for 20 ns without RuBisCO substrates (i.e., 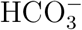, CO_2_, and O_2_) in order to relax the proteins and prepare initial structures used for the US simulations.

Although the central pores were initially dehydrated, they became hydrated in less than 1 ns (Fig. 2). To determine the degree of hydration along each of the pores, a 14-Å radius cylinder covering the entire pore, including its concave and convex funnels, was defined. The z-coordinates of the oxygen atoms of water molecules localized within the cylinder during the last 10 ns of the simulations were recorded and clustered into a histogram with 0.5-Å bins, yielding a distribution profile of the water. The pore radius profiles were calculated using the HOLE program [41] to determine the degree of accessibility of the pores, and the water profiles were normalized with respect to cross-sectional areas along the sections within the pores. The normalized profiles shown in the middle panels of Fig. 2 represent the lowest occupancy site for water molecules at z = 0, which corresponds to the narrowest section, or the bottleneck, of the pore (Fig. 3, right panels). For CsoS1A, this bottleneck is formed by residues G42 and G43. For CcmK4, the equivalent section is formed by residues S41.

**Figure 2:**
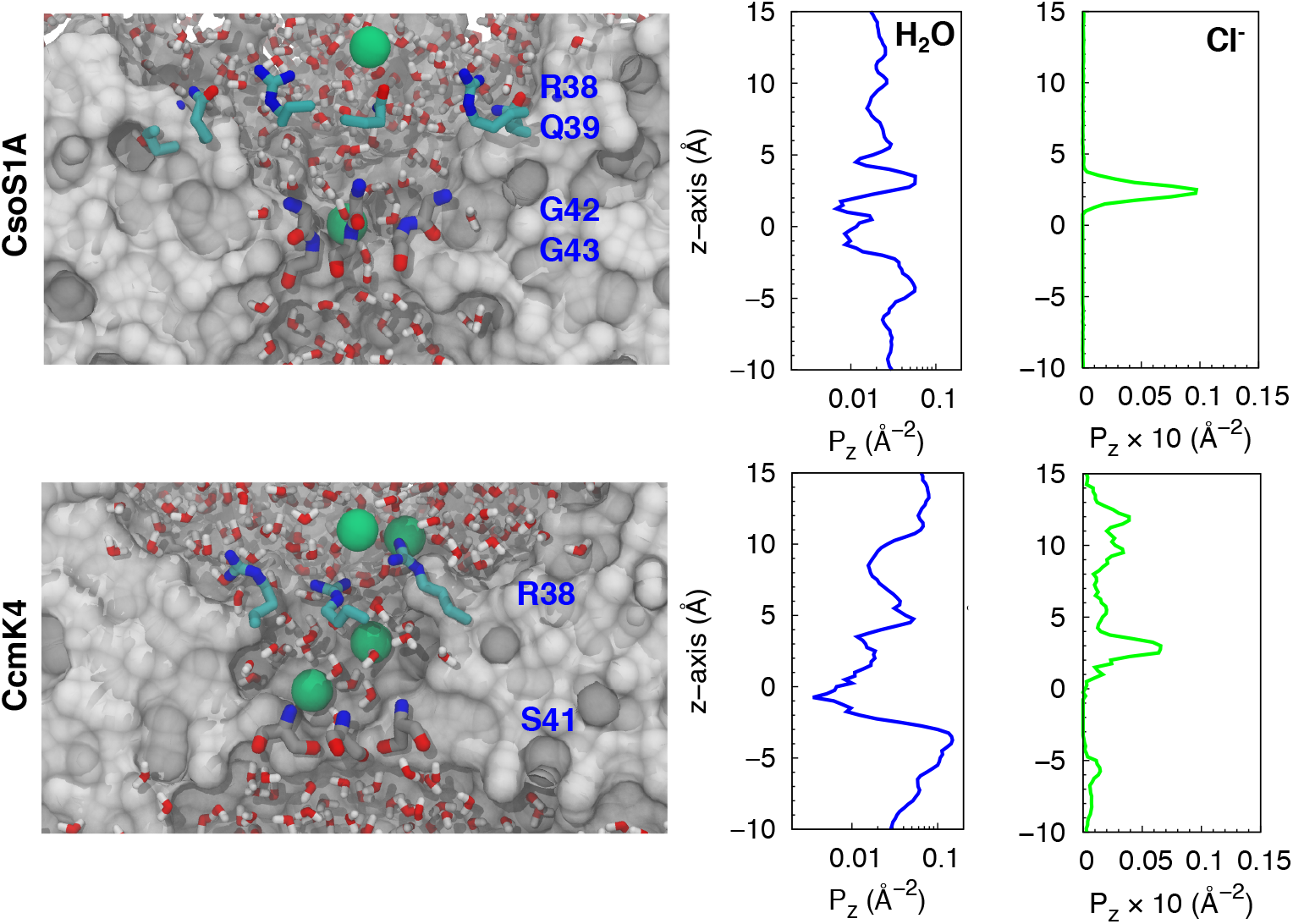
Hydration and Cl^−^ partitioning along the central pores. Left panels, taken from the equilibrium simulations, delineate the localizations of water and Cl^−^ (green spheres) within the central pores. Middle and right panels show the distribution profiles of water molecules and Cl^−^ ions normalized using cross-sectional areas along the sections within the proes. The backbone nitrogen atoms of residues G43 of CsoS1A and S41 of CcmK4 mark the center of the pore and are considered z = 0.

**Figure 3:**
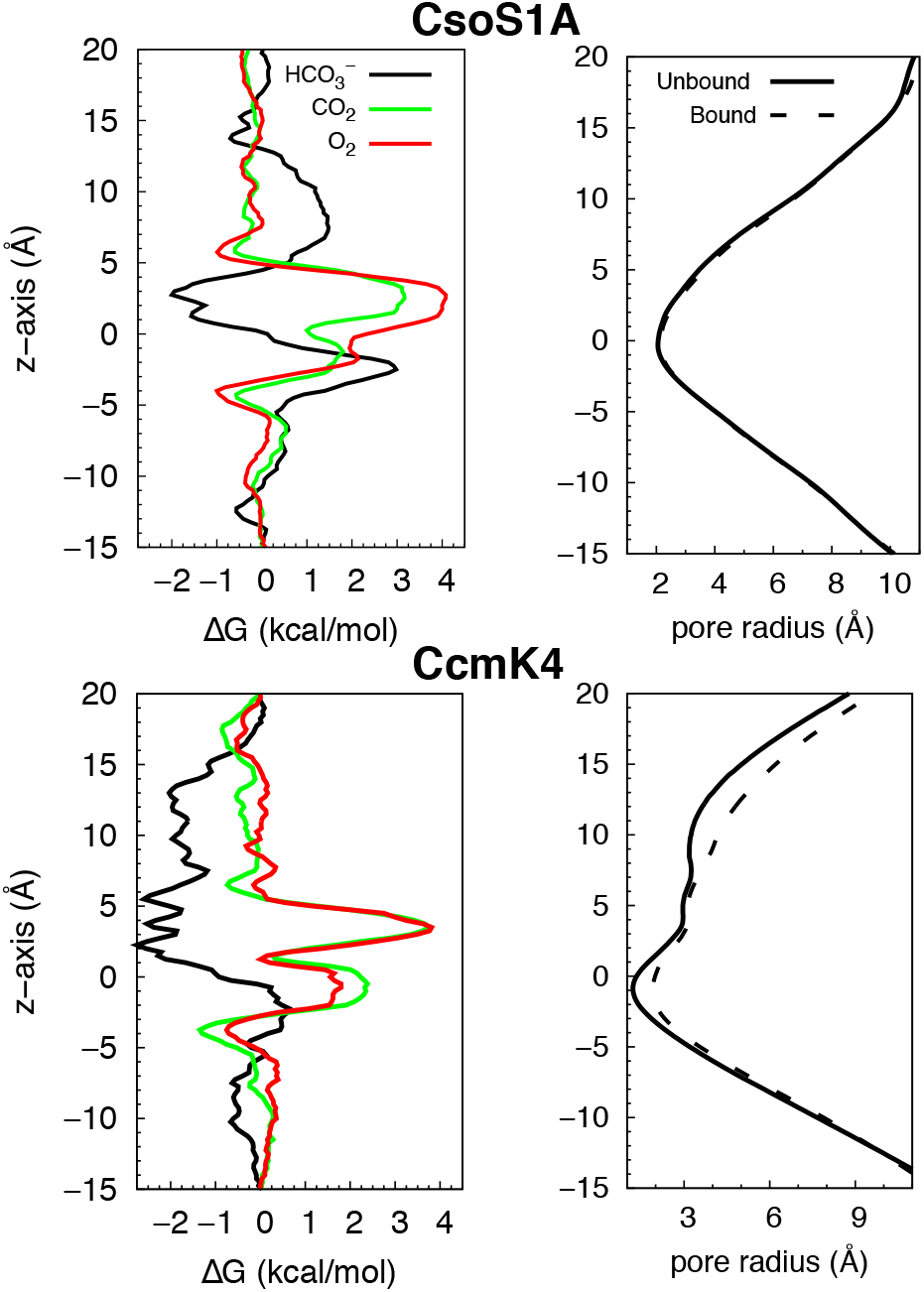
Partitioning of the substrates along the central pores. Left, free energy (ΔG) profiles for inserting substrates of the RuBisCO enzymes along the pores. Right, pore radius profiles of the central pores in the presence and absence of the substrate. 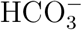 was used to represent the substrate as the other substrates similar profiles showed similar profiles.

These simulations along with previous structural studies suggest favorable anion binding along the central pores. During the 20-ns period of each of the simulations, spontaneous binding of Cl^−^ ions was observed. The Cl^−^ ions bound the amine groups of residues G43 of CsoS1A and those of residues S41 of CcmK4. As shown in the left panels of Fig. 2, the highest Cl^−^ occupancy site was located at z = 3 Å. For CcmK4, apparent accumulation of Cl^−^ ions was also found near residues R38, which lie between z = 8 Å and z = 14 Å. The binding of other ions such as SO_4_^2−^ has also been reported by X-ray crystallography [20, 42]. The crystal structure of CsoS1A, which was used in the simulations, contains one SO_4_^2−^ molecule bound to the amine groups of G43 [20]. The same binding is also found in the crystal structures of CcmK1 and CcmK2, which are homologues of CcmK4 [42]. For CcmK1 and CcmK2, the serine residues equivalent to S41 of CcmK4 bind a SO_4_^2−^ molecule. These results all point to the presence of apositive electrostatic potential within the pore attracting negatively charged molecules.

### Selectivity of anionic substrates

Free energy (ΔG) profiles of 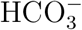, CO_2_, and O_2_ insertion (Fig. 3, left panels) were calculated in order to determine the selectivity of the central pores for these substrates. In agreement with the results of our equilibrium simulations, the resulting free energy profiles demonstrate favorable binding of anionic species (i.e., 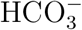) to the central pores, in both CsoS1A and CcmK4. For 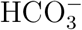 insertion in CsoS1A, a free energy well was found in the region between z = 1 Å and z = 5 Å. The lowest ΔG was located at z = 3 Å and is∼-2 kcal/mol. For CcmK4, favorable 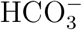 binding regions (relative to bulk water) spanned from z = 1 Å to z = 15 Å, with the global ΔG minimum of ∼-2.5 kcal/mol occurring between z = 2 Å and z = 5 Å. Another binding region with a weaker insertion free energy (∼-2 kcal/mol) is discernible between z = 8 Å and z = 14 Å, and corresponds to the position of a second Cl^−^binding site observed in one of our equilibrium simulations (Fig. 2, bottom panels).

While the binding of 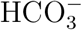 in the central pore is generally favorable, in both CsoS1A and CcmK4 central pores, this substrate has to overcome uphill free energy changes during its translocation from one side of the shell proteins to the other. Both CsoS1A and CcmK4 exhibited high free energy values at z = –2.5 Å (Fig. 3, left panels). These barriers are significantly smaller for 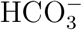 when compared to CO_2_ (see below). Furthermore, the presence of a long attractive region (at 0 *<* z *<* 5 Å) will provide a high probability for the substrate presence right next to the high-energy region.

In contrast to 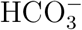, the permeation of CO_2_ and O_2_ through these central pores was more unfavorable with insertion ΔG of 2–4 kcal/mol, extending from z = 5 Å to z = –5 Å. The highest ΔG for CO_2_ and O_2_ insertion, ∼4 kcal/mol, was located at z = ∼3 Å, corresponding to the global ΔG minima for 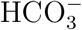 insertion (Fig. 3, left panels). The same section was found to have a high density of water molecules and Cl^−^ ions (Fig. 2). In CsoS1A too, this highest barrier coincides with the region where the mobility of water molecules (as well as other molecular species) was found to be minimum (Fig. 4, left panel).

Because compacted water molecules can hinder the passage of relatively nonpolar CO_2_ and O_2_, we analyzed the mobility of 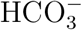, CO_2_ and O_2_ along the central pores by calculating lateral diffusion coefficients (D) of these molecules. The diffusion profiles showed that as any of these molecules approach the bottleneck, their diffusivity becomes significantly diminished, by 10–40 fold (Fig. 4), consistent with a decrease in pore size (Fig. 3, right panels).

**Figure 4:**
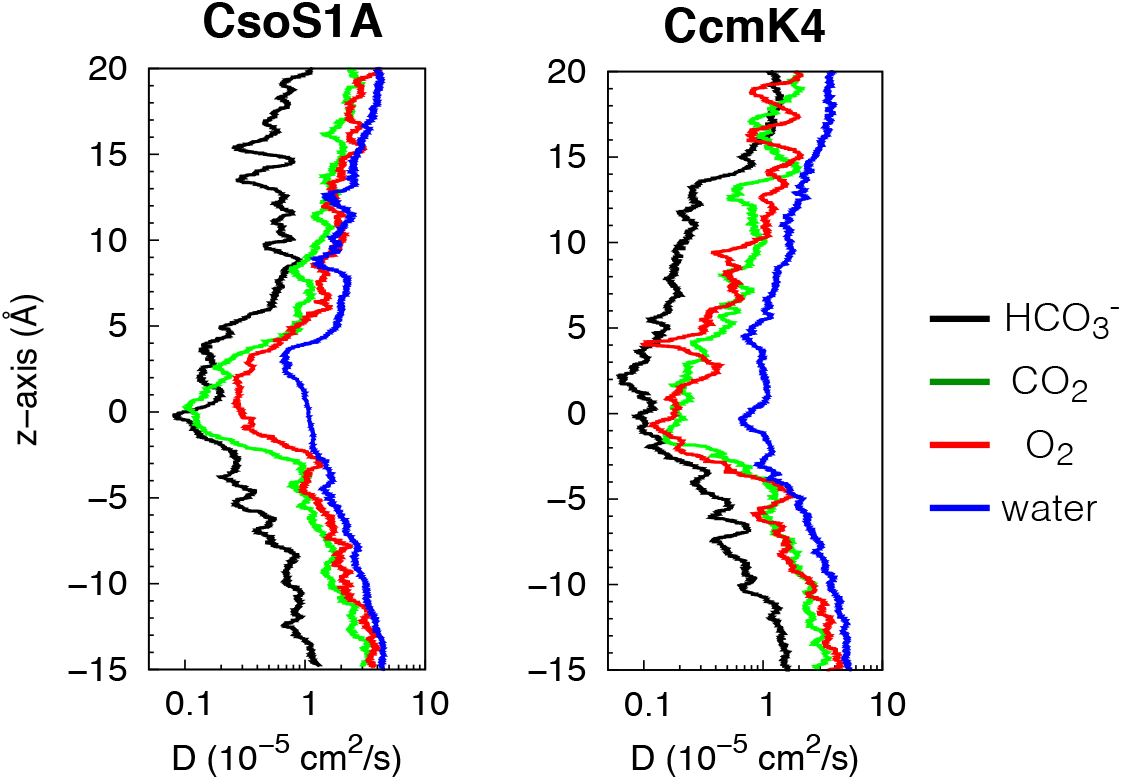
Lateral diffusion coefficient (D) profiles of 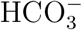, CO_2_, O_2_, and water molecules along the central pores of CsoS1A and CcmK4 hexamers. D is defined as < (Δx) ^2^ + (Δy) ^2^ > / 4Δt, where Δx and Δy were the displacements of the molecule along the x-axis and the y-axis, respectively, at a z position and Δt = 10 ps. < … > denotes mean.

### Dynamics of protein and substrates

The movement of a molecule through a protein may involve not only structural perturbations of the protein, from breathing motions of lining amino acids [43–46] to largescale conformational changes [47–51], but also the dynamics of the passing molecule (substrate or ligand) [52]. Although the pores of CsoS1A and CcmK4 are relatively wide to accommodate a free passage of small molecules, such as 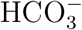, CO_2_ and O_2_, the calculated ΔG profiles indicated that such molecules cannot pass through easily.

Pore radii along the central pores were re-calculated upon the localization of a substrate molecule within the bottleneck. For CsoS1A, the bottleneck pore radius remained ∼2 Å (Fig. 3, upper right panel). For CcmK4, it increased from 1.5 to 2 Å (Fig. 3, lower right panel). This change indicates a slight opening of the pathway, consistent with an increase in the radius of gyration between the six hydroxyl groups of S41 from 3 to 4 Å. In this configuration, the hydroxyl groups of S41 are oriented away from the pathway. Nevertheless, the bottleneck pore radii of both CsoS1A and CcmK4 are ∼2 Å, suggesting that the permeating molecules have to orient themselves just right to transit through the bottleneck.

**Figure 5:**
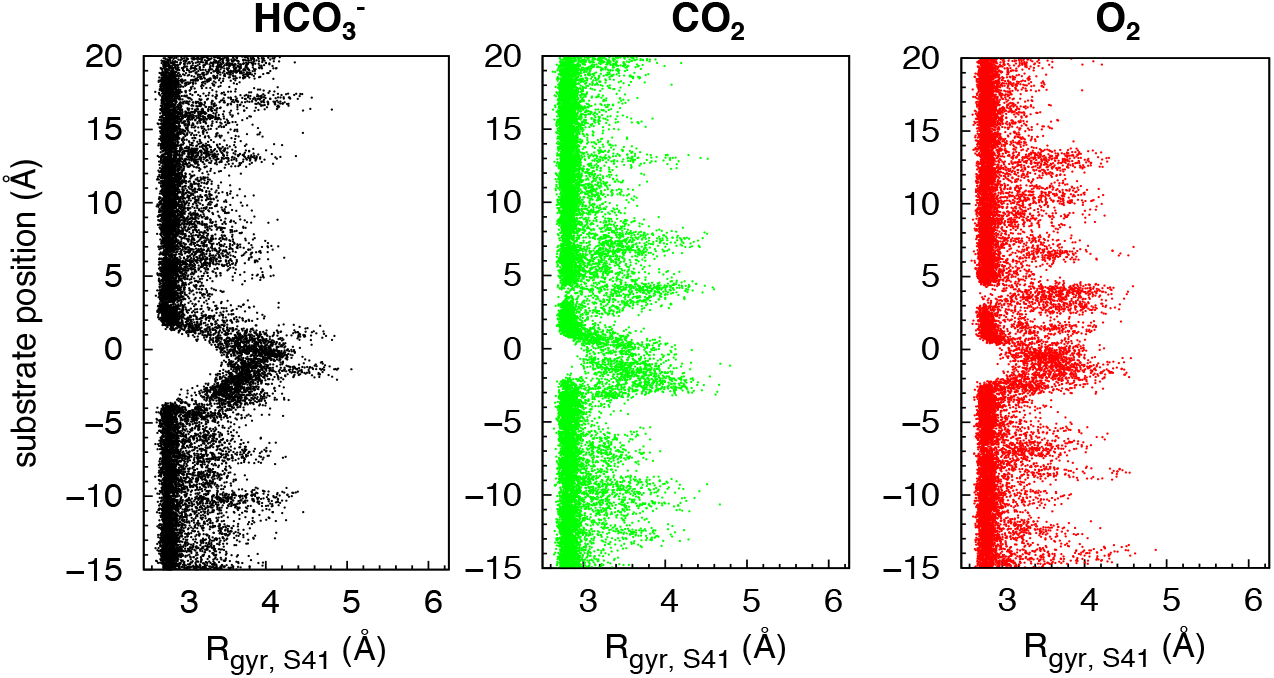
Conformational changes of bottlneck residues in CcmK4 upon substrate binding along the pore determined by calculating the radius of gyration (R_*gyr*_) of the hydroxyl groups of residues S41. This R_*gyr*_was calculated because in the crystal structure, the hydroxyl groups of these bottleneck amino acids appear to constrict the permeation pathway.

Conformational selectivity of the substrates was determined by calculating P_1_ and P_2_ order parameters, shown in Fig. 6, delineating the binding orientation of the substrate molecules upon migrating along the pore. As shown in Fig. 6, the orientation of the substrates was isotropic near the protein surface or in the bulk solution (z<-10 Å or z >15 Å). 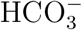 became conformationally restricted when approaching or being within the bottleneck of the central pores. Upon positioning towards the concave surface, its negatively charged carboxylate faced the bottleneck when located at the ΔG minimum or z = 3 Å, as indicated by P_1_ *∼-*1. The molecule flipped by 180° during the transition between the ΔG minimum and the bottleneck, as indicated by P_1_ ∼1. Between the bottleneck and z = –5 Å (convex funnel), it became perpendicularly oriented with respect to the pore, indicated by P_2_ ∼-0.5.

**Figure 6:**
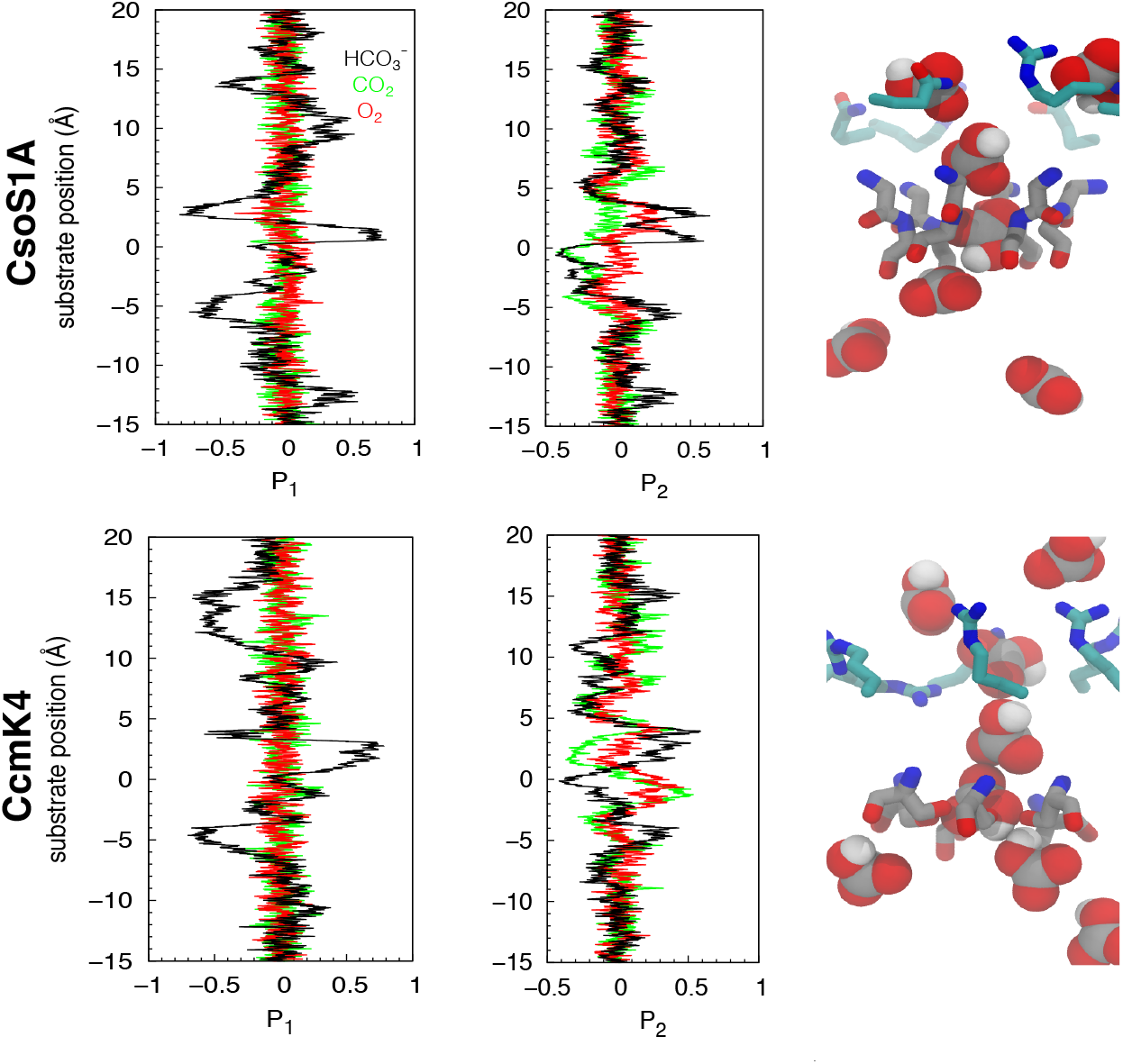
Binding orientation of substrates along the central pores (P_1_ or P_2_ order parameters versus substrate’s positions along the z axis). P_1_ = <cosθ > and P_2_ < <(3cosθ^2^ –1)/2>. P_1_ is ∽±1 when a substrate molecule is oriented parallel to the pore axis and ∼0 when it is oriented either isotropically or perpendicularly with respect to the pore axis. P_2_ differentiates between isotropic average orientation (P_2_ = 0) and orthogonality (P_2_ = –0.5), in which the molecule is in a perpendicular orientation. corresponds to the tilt angle with respect to the pore. For 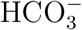,*θ* is the angle between the C-OH bond vector and the pore axis. For linear CO_2_ and O_2_ molecules, *θ* is the angle between the main axis of the substrate and the pore axis. The right panels show several snapshots of 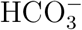 taken from the US simulations.

### Charge distributions along the central pores

Electrostatic potential surface calculations support the preferential selectivity of the central pores of CsoS1A and CcmK4 for anions over nonpolar molecules. Time-averaged electrostatic potentials of the protein complexes were calculated using the structures of the protein hexamers generated from the last 10 ns of the equilibrium simulations (Fig. 7). The calculated electrostatic potential surfaces are consistent with ones previously calculated using the crystal structures [20, 21], supporting the idea that the pores are highly charged.

These calculations also rationalize the features observed in the calculated ΔG profiles (Fig. 3, left panels). For CcmK4, the central pore is positive, which provides attraction for 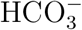 molecules. For CsoS1A, on the other hand, electrostatic potentials along the pore were asymmetrically distributed. The funnel from the concave surface to the bottleneck was positively charged, whereas the one from its convex surface was negatively charged (Fig. 7). This charge separation provides an explanation for the nature of the permeation ΔG barrier for 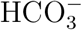 at the bottleneck. Because the convex surface is negatively charged, it is unfavorable for negatively charged molecules, such as 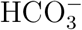, to spontaneously transit through the bottleneck (z = –2.5 Å) from the concave surface to the convex surface. This is in agreement with the ΔG calculations, which showed a 2 kcal/mol higher barrier for a 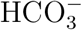 molecule to enter the pore from the concave surface than from the convex surface.

**Figure 7:**
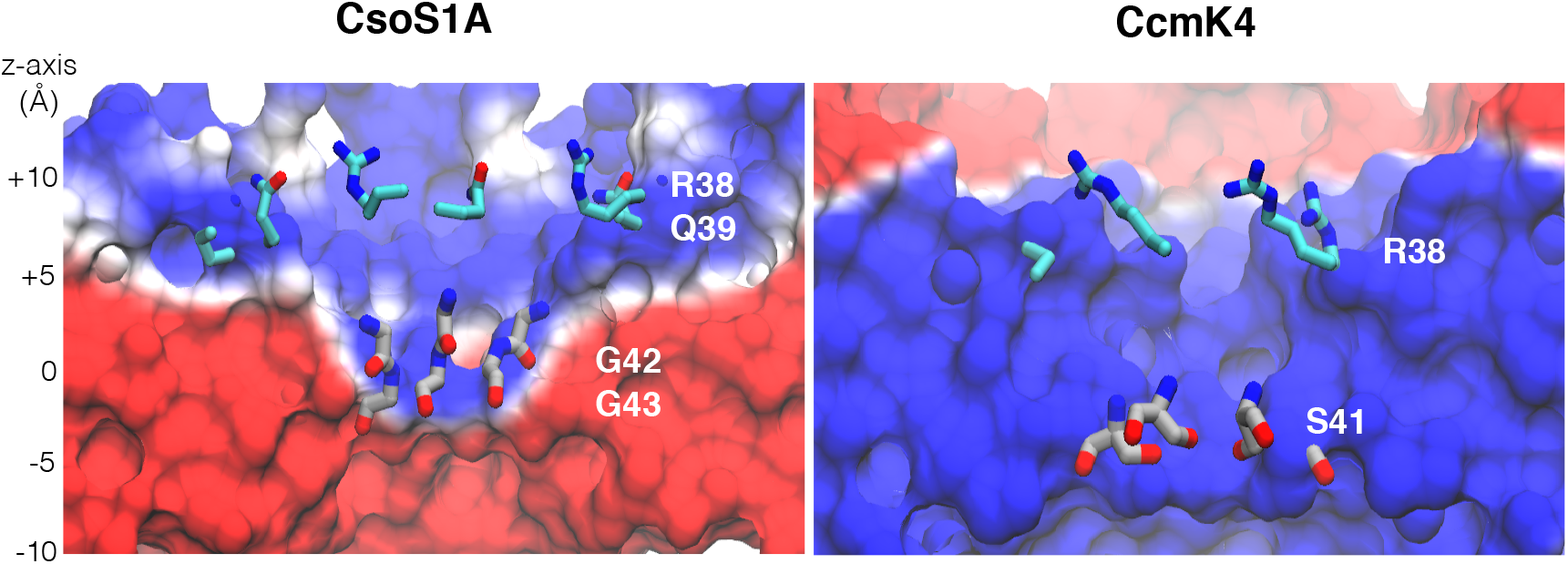
Electrostatic potential maps of the shell proteins. The maps were calculated using PMEpot plugin of VMD [24, 53], which uses the particle mesh Ewald method [54]. The ensemble of protein conformations collected in every 10 ps was used in each calculation. Electrostatic potentials shown in this figure are with threshold ranging from –20 (red) to +20 (blue) kT/e. “positive” z is towards the concave surface, while “negative” z is towards the convex surface.

## Conclusion

The results of the simulations and free energy calculations presented in this study show that the central pores of carboxysome shell protein complexes favor 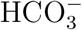 over either CO_2_ or O_2_. Once within the carboxysomal lumen, 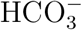 is converted to CO_2_ by carbonic anhydrase. The poor permeability of the carboxysome shell to CO_2_ and O_2_ can improve the productivity of RuBisCO in two ways; not only can it minimize outward leakage of CO_2_ from carboxysomal lumen upon production by carbonic anhydrase, but it will also prevent unwanted entry of O_2_ into the lumen. These results substantiate, at a molecular level, how carboxysomes maintain the high local CO_2_ concentrations around the RuBisCO enzymes necessary for adequate performance of these otherwise inefficient enzymes. Consistent with the idea that the carboxysome shell acts as a barrier against the permeation of CO_2_, and that of O_2_, the experimental study by Dou et al [55] on *H. neapolitanus* suggests that the CO_2_ supply into the carboxysome is provided mainly thorugh the entry of 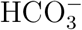 molecules. The activity of freely soluble RuBisCO enzymes was compared to the activity of the enzymes contained inside the intact carboxysome. The intact carboxysome devoid of carbonic anhydrase showed a 3-fold increase in *K_m_* of CO_2_ with no change in *V_max_*, while the ruptured carboxysome and carboxysome-free RuBisCO enzymes showed similar *K_m_* and *V_max_* to the wild type. As a consequence, they found that *H. neapolitanus* mutants lacking carbonic anhydrase required elevated CO_2_ to grow.

The observed substrate selectivity appears to originate from electrostatic properties of the central pores. Positive electrostatic potentials along the central pores establish strong binding affinities for negatively charged molecules, such as 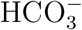 and Cl^−^ ions. This notion is supported by the calculated favorable insertion free energies of 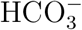 from the umbrella sampling simulations and the captures of spontaneous Cl^−^ binding during the equilibrium simulations. Favorable binding of 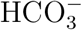 facilitates its passage through the pore.

## Acknowledgements

We thank Dr. Catherine Oikonomou for help revising the manuscript. This work was supported in part by the National Institutes of Health (NIH P41-GM104601, U01-GM111251 and U54-GM087519 to E.T.), the Office of Naval Research (ONR N00014–16-1–2535 to E.T.), and the Howard Hughes Medical Institute (to G.J.J.). P.M. gratefully acknowledges a previous support as a trainee of the Molecular Biophysics Training Program by the NIH (T32-GM008276) during his graduate study.

